# Paragraph: A graph-based structural variant genotyper for short-read sequence data

**DOI:** 10.1101/635011

**Authors:** Sai Chen, Peter Krusche, Egor Dolzhenko, Rachel M. Sherman, Roman Petrovski, Felix Schlesinger, Melanie Kirsche, David R. Bentley, Michael C. Schatz, Fritz J. Sedlazeck, Michael A. Eberle

## Abstract

Accurate detection and genotyping of structural variations (SVs) from short-read data is a long-standing area of development in genomics research and clinical sequencing pipelines. We introduce Paragraph, an accurate genotyper that models SVs using sequence graphs and SV annotations. We demonstrate the accuracy of Paragraph on whole-genome sequence data from three samples using long read SV calls as the truth set, and then apply Paragraph at scale to a cohort of 100 short-read sequenced samples of diverse ancestry. Our analysis shows that Paragraph has better accuracy than other existing genotypers and can be applied to population-scale studies.

## Background

Structural variants (SVs) contribute to a large fraction of genomic variation and have long been implicated in phenotypic diversity and human disease^1–3^. Whole-genome sequencing (WGS) is a common approach to profile genomic variation, but compared to small variants, accurate detection and genotyping of SVs still remains a challenge^4,5^. This is especially problematic for a large number of SVs that are longer than the read lengths of short-read (100-150 bp) high-throughput sequence data, as a significant fraction of SVs has complex structures that can cause artifacts in read mapping and make it difficult to reconstruct the alternative haplotypes^6,7^.

Recent advances in long read sequencing technologies, (e.g. Pacific Biosciences and Oxford Nanopore Technologies), have made it easier to detect SVs, including those in low complexity and non-unique regions of the genome. This is chiefly because, compared to short reads, long (10-50kbp) reads can be more reliably mapped to such regions and are more likely to span entire SVs^8–10^. These technologies combined with data generated by population studies using multiple sequencing platforms, are leading to a rapid and ongoing expansion of the reference SV databases in a variety of species^11–13^.

Currently, most SV algorithms analyze each sample independent of any prior information about the variation landscape. The increasing availability and completeness of a reference database of known SVs, established through long read sequencing and deep coverage short-read sequencing, makes it possible to develop methods that use prior knowledge to genotype these variants. Furthermore, if the sequence data remains available they can be re-genotyped using new information as the reference databases are updated. Though the discovery of *de novo* germline or somatic variants will not be amenable to a genotyping approach, population studies that involve detection of common or other previously known variants will be greatly enhanced by genotyping using a reference database that is continually updated with newly discovered variants.

Targeted genotyping of SVs using short-read sequencing data still remains an open problem^14^. Most targeted methods for genotyping are integrated with particular discovery algorithms and require the input SVs to be originally discovered by the designated SV caller^15–17^, require a complete genome-wide realignment^18,19^ or need to be optimized on a set of training samples^12,20^. In addition, insertions are generally more difficult to detect than deletions using short-read technology, and thus are usually genotyped with lower accuracy or are completely excluded by these methods^21–23^. Finally, consistently genotyping SVs across many individuals is difficult because most existing genotypers only support single-sample SV calling.

Here, we present a graph-based genotyper, Paragraph, that is capable of genotyping SVs in a large population of samples sequenced with short reads. The use of a graph for each variant makes it possible to systematically evaluate how reads align across breakpoints of the candidate variant. Paragraph can be universally applied to genotype insertions and deletions represented in a variant call format (VCF) file, independent of how they were initially discovered. This is in contrast to many existing genotypers that require the input SV to have a specific format or to include additional information produced by a specific *de novo* caller^14^. Furthermore, compared to alternate linear-reference based methods, the sequence graph approach minimizes the reference allele bias and enables the representation of pan-genome reference structures (e.g. small variants in the vicinity of an SV) so that variants can be accurate even when variants are clustered together^24–27^.

We compare Paragraph to five popular SV detection and genotyping methods and show that the performance of Paragraph is an improvement in accuracy over the other methods tested. Our test set includes 20,385 SVs (9,287 deletions and 11,117 insertions) across three human samples for a total of 60,389 genotypes (38,265 alternative and 22,124 homozygous reference genotypes). Against this test set, Paragraph achieves a recall of 0.86 and a precision of 0.91. By comparison the most comprehensive alternative genotyping method we tested achieved 0.76 recall and 0.85 precision across deletions only. In addition, the only discovery-based SV caller we tested that could identify both insertions and deletions, had a recall of 0.35 for insertions compared to 0.88 for Paragraph. Finally, we showcase the capability of Paragraph to genotype on a population-scale using 100 deep-coverage WGS samples, from which we detected signatures of purifying selection of SVs in functional genomic elements. Combined with a growing and improving catalog of population-level SVs, Paragraph will deliver more complete SV calls and also allow researchers to revisit and improve the SV calls on historical sequence data.

## Result

### Graph-based genotyping of structural variations

For each SV defined in an input variant call format (VCF) file, Paragraph constructs a directed acyclic graph containing paths representing the reference sequence and possible alternative alleles (**Figure 1**) for each region where a variant is reported. Each node represents a sequence that is at least one nucleotide long. Directed edges define how the node sequences can be connected to form complete haplotypes. The sequence for each node can be specified explicitly or retrieved from the reference genome. In the sequence graph, a branch is equivalent to a variant breakpoint in a linear reference. In Paragraph, these breakpoints are genotyped independently and the genotype of the variant can be inferred from genotypes of individual breakpoints (see **Methods**). Besides genotypes, several graph alignment summary statistics, such as coverage and mismatch rate, are also computed which are used to assess quality, filter and combine breakpoint genotypes into the final variant genotype. Genotyping details are described in the **Methods** section.

**Figure 1.**
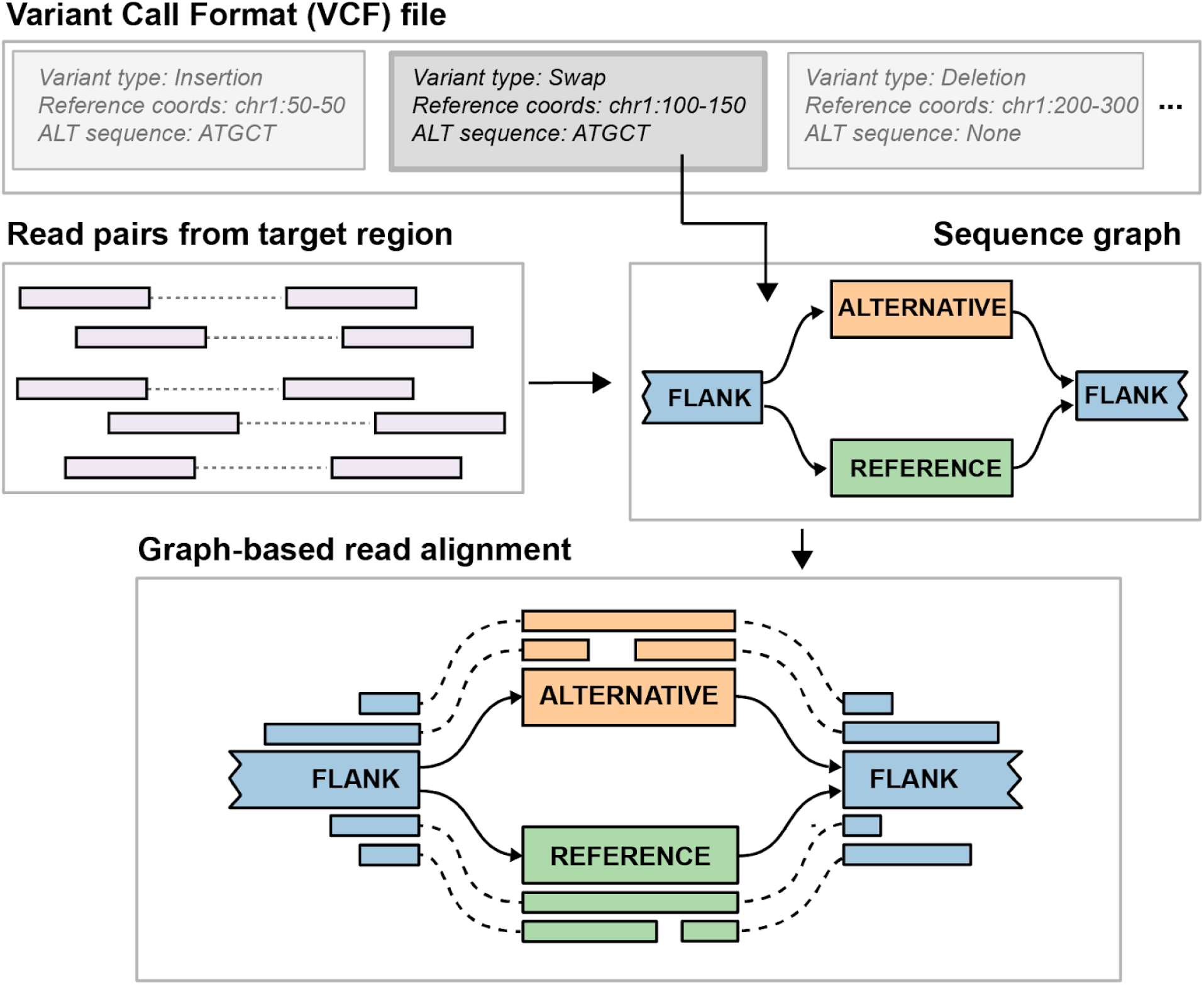
Overview of the SV genotyping workflow implemented in Paragraph. The illustration shows the process to genotype a blockwise sequence swap. Starting from an entry in a VCF file that specifies the SV breakpoints and alternative allele sequences, Paragraph constructs a sequence graph containing all alleles as paths of the graph. Colored rectangles labeled FLANK, ALTERNATIVE and REFERENCE are nodes with actual sequences and solid arrows connect these nodes are edges of the graph. All reads from the original, linear alignments that aligned near or across the breakpoints are then realigned to the constructed graph. Based on alignments of these reads, the SV is genotyped as described in the **Methods** section.

### Construction of a long read-based ground truth

To estimate the performance of Paragraph and other existing methods, we built a long read ground truth (LRGT) from SVs called in three samples included in the Genome in a Bottle (GIAB)^11,28^ project data: NA12878 (HG001), NA24385 (HG002) and NA24631 (HG005). Long read data from these three individuals was generated on a Pacific Biosciences (PacBio) Sequel system using the Circular Consensus Sequencing (CCS) technology (sometimes called “HiFi” reads)^29^. Each sample was sequenced to an average of 30 fold depth and ∼11,100 bp read length. Previous evaluations showed high recall (0.91) and precision (0.94) for SVs called from PacBio CCS NA24385 with similar coverage levels against the GIAB benchmark dataset in confident regions^11,29^. Thus, indicating SVs called from CCS data can be effectively used as ground truth to evaluate the performance of SV genotypers and callers.

For each sample, we called SVs (50bp+) as described in the **Methods** and identified a total of 65,108 SV calls (an average 21,702 SVs per sample) representing 38,709 unique autosomal SVs. In addition, we parsed out SV loci according to regions with a single SV across the samples and those with multiple different SVs and identified that 38,239 (59%) of our SV calls occur as single, unique events in the respective region and the rest 26,869 (41%) occur in regions with one or more nearby SVs (**Figure S1**). Recent evidence suggests that a significant fraction of novel SVs could be tandem repeats with variable lengths across the population^30,31^ and we found that 49% of the singleton unique SVs are completely within the UCSC Genome Browser Tandem Repeat (TR) tracks while 93% of the clustered unique SVs are within TR tracks. Because regions with multiple variants will pose additional complexities for SV genotyping that are beyond the scope of the current version of Paragraph, we limited our LRGT to the 9,238 deletions and 10,870 insertions that are not confounded by the presence of a different nearby or overlapping SV (see **Methods**). Considering all three samples, there are: 1) 4,260/4,439 deletions/insertions that occurred in just one sample, 2) 2,258/2,429 deletions/insertions that occurred in two samples and 3) 2,720/4,002 deletions/insertions that occurred in all three samples. With short-read sequencing also available for these three samples, we are able to test any SV genotyping method and can estimate recall and precision using the long read genotypes as the ground truth.

### Test for recall and precision

To evaluate the performance of different methods, we genotyped the LRGT SVs on short-read data of NA12878 (63x), NA24385 (35x) and NA24631 (40x) using Paragraph and two widely-used SV genotypers, SVTyper^16^ and Delly Genotyper^17^. Additionally, we ran three methods that independently discover SVs (i.e. *de novo* callers), Manta^21^, Lumpy^32^ and Delly^17^. Because the genotyping accuracy of classifying homozygous versus heterozygous alleles may vary for the short and long-read methods used here, we focus our test on the presence/absence of variants and not genotyping concordance. Thus, we define a variant as a true positive (TP) if LRGT also has a call in the same sample and a false positive (FP) if LRGT did not call a variant in that sample. We have 38,239 individual alternative genotypes in LRGT to calculate TPs and 22,085 individual reference genotypes in LRGT to calculate FPs. Since some of the methods are not able to call certain sizes or types of SVs we only tested these methods on a subset of the SVs when calculating recall and precision.

Paragraph has the highest recall: 0.84 for deletions and 0.88 for insertions (**Table 1**) among all the genotypers and *de novo* callers tested. Of the genotypers, Paragraph had the highest genotype concordance compared to the LRGT genotypes (**Table S1**). The precision of Paragraph is estimated as 0.92 for deletions, which is 7% higher than Delly Genotyper (0.85), and 0.89 for insertions. Though SVTyper had the highest precision (0.98) of all the methods tested it achieved that by sacrificing recall (0.70). Furthermore, SVTyper is limited to deletions longer than 100 bp. When measuring precision only on 100bp+ deletions, Paragraph has a slightly lower precision (0.93) than SVTyper (0.98) but the recall is 12% higher (0.82 vs SVTyper 0.70). Combining recall and precision, Paragraph has the highest F-score among all genotypers also for this subset of 100bp+ deletions (0.88 vs 0.80 for Delly Genotyper and 0.82 for SVTyper). In addition, we tested another short-read genotyper, BayesTyper, a kmer-based method and estimated a recall of 0.47 and precision of 0.94 across all of the LRGT SVs. The low recall of BayesTyper is because it produced no genotype call for 56% of the LRGT SVs. We speculate that this may be largely caused by sequencing errors that would have a greater impact on methods that require exact matches of kmers.

**Table 1.** Performance of different genotypers and *de novo* callers, measured against 50bp or longer SV from our LRGT. Genotyping/calling was evaluated on short read data of the three samples sequenced with 150 bp paired-end reads on Illumina platforms. As SVTyper and Lumpy are limited to deletions longer than 100bp, they have fewer tested SVs than other methods.

| Type | Deletion |  |  |  |  |  | Insertion |  |
| --- | --- | --- | --- | --- | --- | --- | --- | --- |
|  | Paragraph | Delly Genotyper | SVTyper (100+ bp) | Manta | Delly | Lumpy (100+ bp) | Paragraph | Manta |
| #Tested TPs | 16,936 | 16,936 | 11,160 | 16,936 | 16,936 | 11,160 | 21,303 | 21,303 |
| Recall | 0.84 | 0.76 | 0.70 | 0.62 | 0.61 | 0.64 | 0.88 | 0.35 |
| #Tested FPs | 10,778 | 10,778 | 6,960 | - | - | - | 11,307 | - |
| Precision | 0.92 | 0.85 | 0.98 | - | - | - | 0.89 | - |
| F-score | 0.88 | 0.80 | 0.82 | - | - | - | 0.88 | - |

Since genotyping performance is often associated with SV length (e.g. depth-based genotypers usually perform better on larger SVs than smaller ones), and some of the tested methods only work for SVs above certain deletion/insertion sizes, we partitioned the LRGT SVs by length and further examined the recall of each method (**Figure 2**). In general, for deletions between 50bp and ∼1,000bp, the genotypers (Paragraph, SVTyper, and Delly Genotyper) have better recall than the *de novo* callers (Manta, Lumpy, and Delly). SVTyper and Paragraph have comparable recall for larger (>300bp) deletions, and in that size range, Delly Genotyper has lower recall than these two. For smaller deletions (50-300 bp), the recall for Paragraph (0.83) remains high while we observe a slight drop in the recall of Delly Genotyper (0.75) and a larger drop in the recall of SVTyper (0.43). We speculate that this is because SVTyper mainly relies on paired-end (PE) and read-depth (RD) information and will therefore be less sensitive for smaller events. Only Paragraph and Manta were able to call insertions and while Paragraph (0.88) has consistently high recall across all insertion lengths, Manta (0.35) has a much lower recall which drops further for larger insertions.

**Figure 2.**
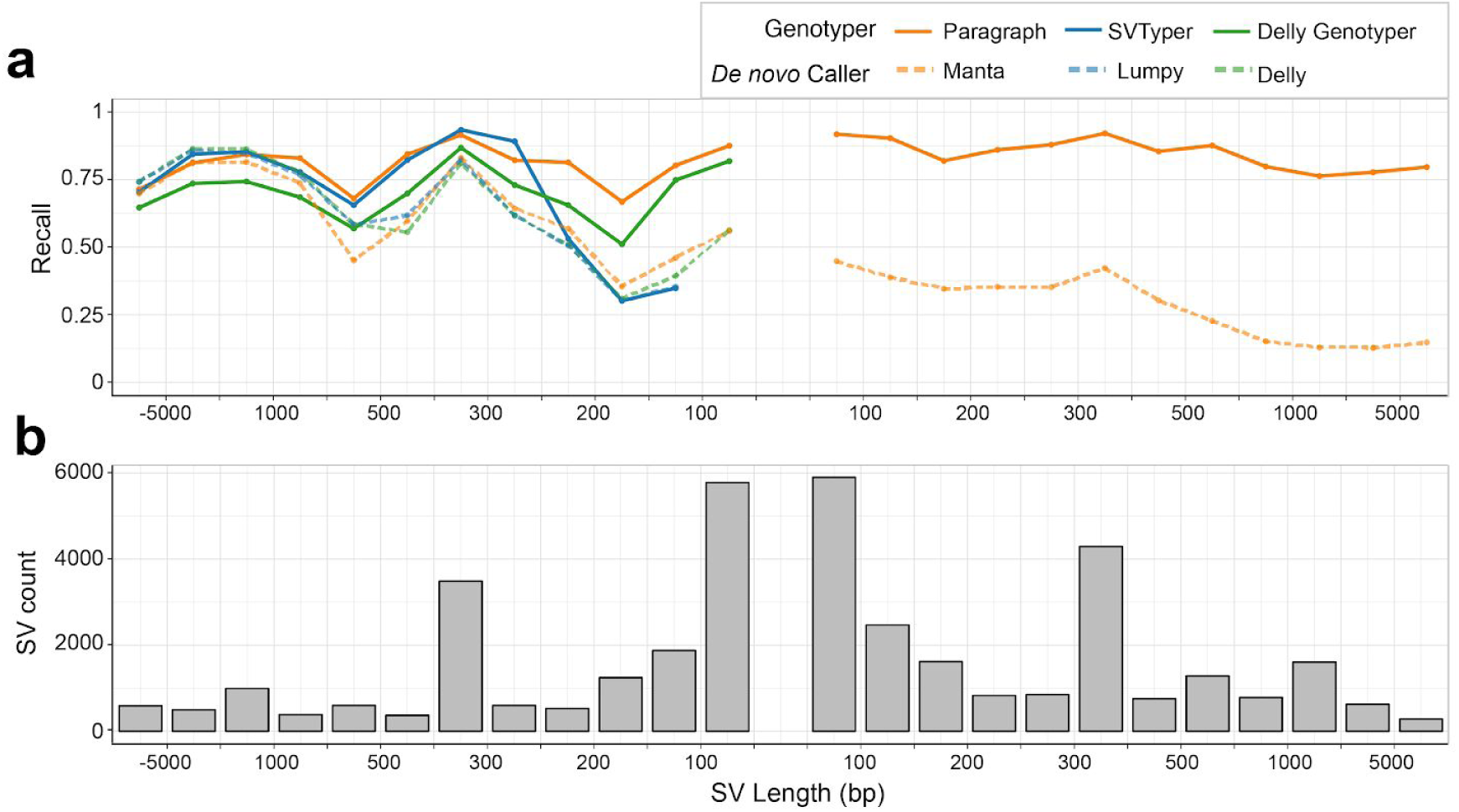
Estimated recall of different methods, partitioned by SV length. Recall was estimated on the three samples using LRGT as the truth set. A negative SV length indicates a deletion and a positive SV length indicates an insertion. Colored lines in (a) show recall of different methods; Solid grey bars in (b) represent the count of SVs in each size range in LRGT. The center of the plot is empty since SVs must be at least 50 bp in length.

We additionally partitioned the precision of each genotyper by SV length (**Figure S2**). The result suggests that false positives are more likely to occur in small SVs than in large ones. Paragraph has a consistent precision for deletions and insertions, while the only comparable method in genotyping very small deletions (50-100bp), Delly Genotyper, has a precision drop in this range (**Figure S2**). We further examined Paragraph FPs in one of the tested samples, NA24385 and found nearly all of the FP deletions (91%) and the FP insertions (90%) are completely within TR regions. We performed a visual inspection of the 21 FP deletions and 83 FP insertions that are outside of TRs: 12% (12) have two or more supporting reads for an SV but were not called by the long read caller in LRGT; 40% (42) have one or more large indels (longer than 10bp) in the target region; 48% (50) have no evidence of variants in the long read alignments in the target region and thus these FPs are likely to come from short-read alignment artifacts.

So far, we tested the recall using high depth data (>35x) with 150bp reads but some studies may use shorter reads and/or lower read depths. To quantify how either shorter reads or lower depth will impact genotyping performance, we evaluated data of different read lengths and depths by downsampling and trimming reads from our short-read data of NA24385. Generally, shorter read lengths are detrimental to recall; reductions in depth have less of a deleterious effect until the depth is below ∼20x (**Figure S3**).

### Genotyping with breakpoint deviations

The LRGT data we used here will be both costly and time-consuming to generate in the near term because generating long read CCS data is still a relatively slow and expensive process. An alternative approach to build up a reference SV catalog would be to sequence many samples (possibly at lower depth) using PacBio contiguous long reads (CLR) or Oxford Nanopore long reads rather than CCS technology and derive consensus calls across multiple samples. The high error rates (∼10-15%) of these long reads may result in errors in SV descriptions especially in low-complexity regions where just a few errors in the reads could alter how the reads align to the reference. Since Paragraph realigns reads to a sequence graph using stringent parameters, inaccuracies in the breakpoints may result in a decreased recall.

To understand how the genotypers perform with input SVs that have imprecise breakpoints, we called SVs from CLR data of NA24385 that were generated on a PacBio RS II platform. 9,534 out of the total 12,776 NA24385 SVs in LRGT closely match those generated from the CLR data (see **Methods** for matching details). Of these, 658 (17%) deletions and 806 (14%) insertions have identical breakpoints in the CLR and CCS SV calls. The remaining 3,306 deletions and 4,763 insertions, although in approximately similar locations, have differences in representations (breakpoints and/or insertion sequences). Assuming breakpoints found using the CCS data within the LRGT SVs are correct, we consider deviations in the CLR breakpoints as errors in this sample. For the matching deletions between LRGT and CLR calls but with deviating breakpoints, Paragraph recall decreased from 0.97 to 0.83 when genotyped the CLR-defined deletions. Overall, there is a negative correlation between Paragraph recall and breakpoint deviations: the larger the deviation, the less likely the variant can be genotyped correctly (**Figure 3**). While deviations of a few base pairs can generally be tolerated without issue, deviations of 20 bp or more reduce recall to around 0.44. For insertions with differences in breakpoints and/or insertion sequences, Paragraph recall decreased from 0.88 to 0.66 when genotyped the CLR-defined insertions. We also investigated how inaccurate breakpoints impact insertion genotyping, but found no clear trend between recall and base-pair deviation in breakpoints.

**Figure 3.**
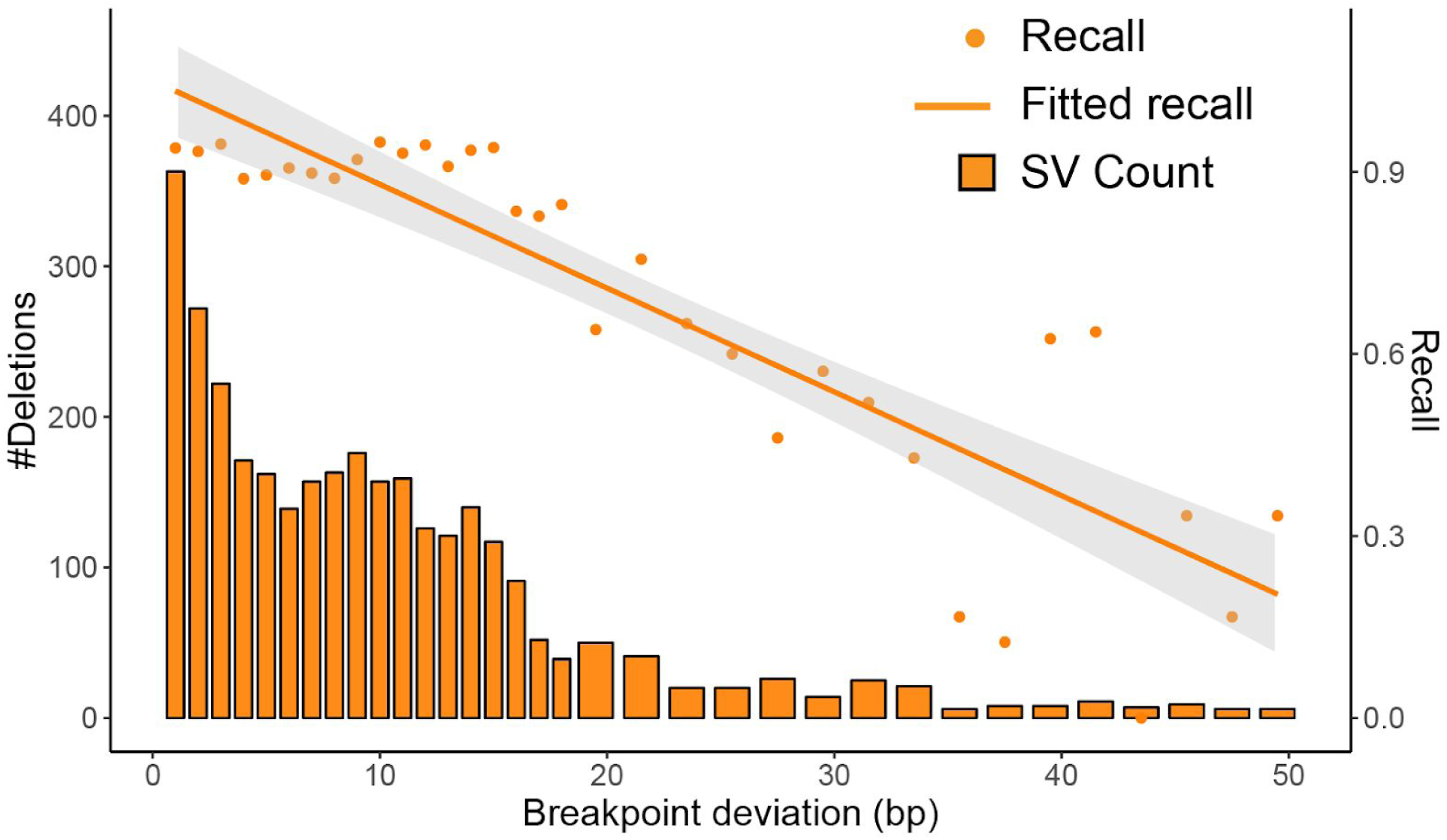
Demonstration of the impact of recall when tested SVs include errors in their breakpoints. Breakpoint deviations measure the differences in positions between matching deletions in the CLR calls and in LRGT. Paragraph recall was estimated using CLR calls as genotyping input and TPs in LRGT as the ground truth. Breakpoint deviations were binned at 1bp for deviations less than 18bp and at 2bp for deviations larger or equal to 19bp. Solid bars show the number of deletions in each size range (left axis). Points and the solid line shows the recall for individual size and the overall regression curve (right axis).

On the same set of CLR calls, we estimated the impact of breakpoint deviation on SVTyper and Delly Genotyper (**Figure S4**). Similar to Paragraph, the split-read genotyper, Delly Genotyper, shows the same negative relationship between its recall and breakpoint deviations. As a contrast, SVTyper, which genotype SVs mostly using information from read depth and pair-read insert size distribution, does not depend much on breakpoint accuracy and is not significantly affected by deviations in breakpoints.

### Genotyping in tandem repeats

We identified that most of the SVs having breakpoint deviations between the CLR calls and LRGT are in low complexity regions: of the 8,069 matching SVs with breakpoint deviations, 3,217 (77%) are within in TRs. SVs within TRs have larger breakpoint deviations in CLR calls from the true breakpoints than those not in TRs: 35% of the SVs with smaller (<=10 bp) deviations are within TRs while 66% of the SVs with larger breakpoint deviations (>20 bp) are within TRs. Additionally, we found that 59% of the FNs and 77% of the FPs in NA24385 occur in SVs that are completely within TRs. To further understand the impact of TRs on the performance of Paragraph, we grouped LRGT SVs according to whether they are in TRs and plotted Paragraph recall binned by SV lengths. Paragraph has a better recall in SVs that are outside of TRs (0.89 for deletions and 0.90 for insertions), compared to its recall in SVs that are within TRs (0.74 for deletions and 0.83 for insertions) (**Figure 4a**). Small (<200bp) SVs are much more likely to be within TR (∼75%) than large (>1,000bp) SVs (∼35%) (**Figure 4b**), and that matches our earlier observation that Paragraph and other genotypers have decreased recall and precision, in small SVs.

**Figure 4.**
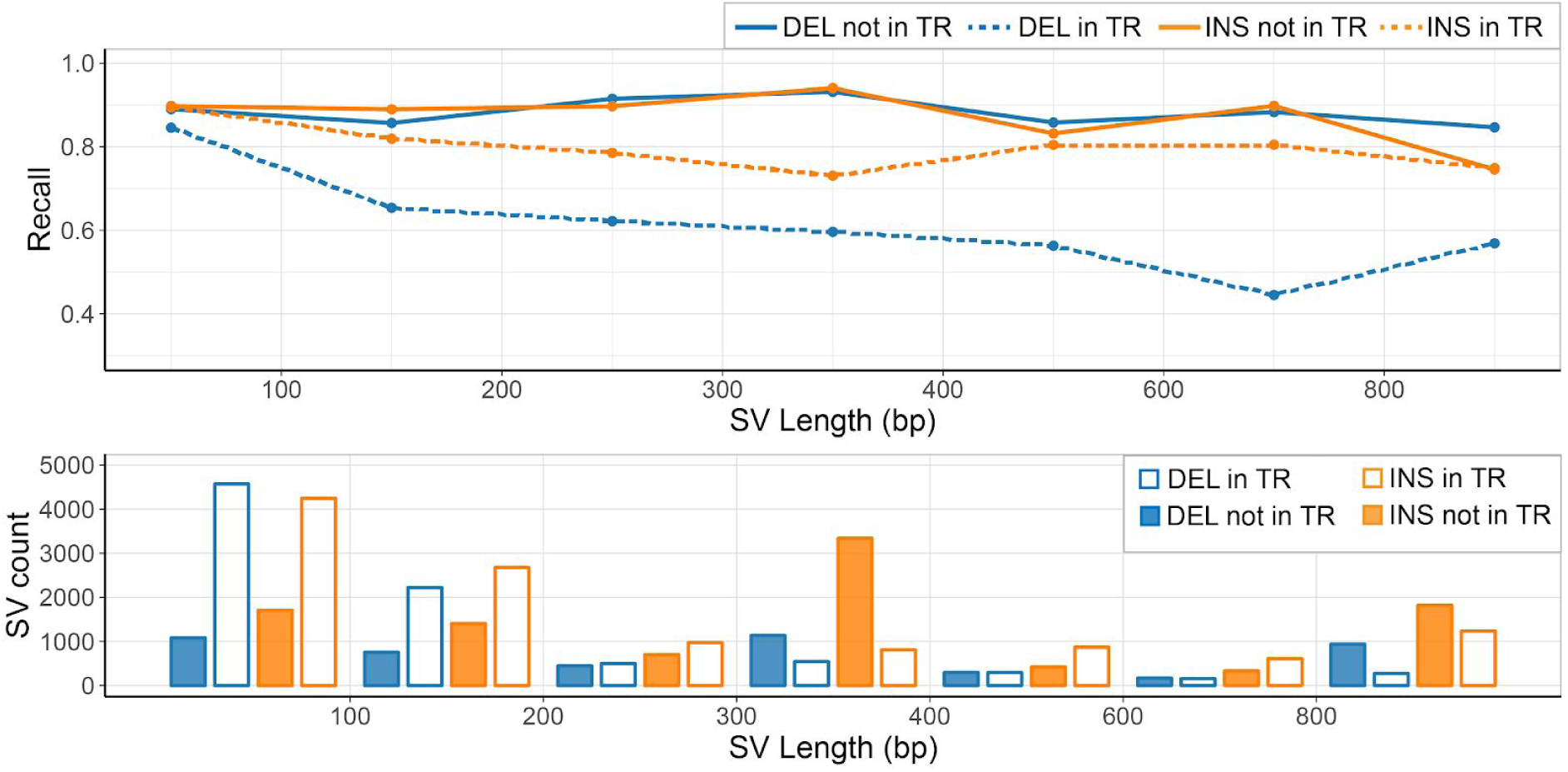
The impact of TRs on SV recall. a) Estimated Paragraph recall from LRGT, partitioned by SV length and grouped by their positioning with TRs. b) LRGT SV count partitioned by length and grouped by their positionings with TRs.

When building our LRGT, we excluded SVs with other nearby SVs in one or more samples (named as clustered SVs in **Construction of long read-based ground truth** section). The majority of these SVs (93%) are within TRs, therefore benchmarking against these clustered SVs could be informative to quantify the impact of TRs in SV genotyping. As none of the tested methods could model each SV cluster as a whole without an appropriate annotation, we instead model each of the SVs in the clusters as a single SV and evaluated the performance of Paragraph and other methods on the same three samples using long read genotypes of these clustered SVs as the underlying truth (**Table S2**). All methods have a lower recall and precision in the clustered SVs than in LRGT highlighted by their reduced F-scores: Paragraph (0.64 vs. 0.88), Delly Genotyper (0.58 vs. 0.80) and SVTyper (0.42 vs. 0.82). The three *de novo* callers have a deletion recall of 0.15-0.20 in the clustered SVs, much lower than their recall of 0.61-0.64 in LRGT.

### Population-scale genotyping across 100 diverse human genomes

A likely use case for Paragraph will be to genotype SVs from a reference catalog for more accurate assessment in a population or association studies. To further test and demonstrate Paragraph in this application, we genotyped our LRGT SVs in 100 unrelated individuals (not including NA24385, NA12878 or NA24631) from the publicly-available Polaris sequencing resource (https://github.com/Illumina/Polaris). This resource consists of a mixed population of 46 Africans (AFR), 34 East Asians (EAS) and 20 Europeans (EUR). All of these samples were sequenced on Illumina HiSeq X platforms with 150 bp paired-end reads to at least 30-fold depth per sample.

Most deletions occur at a low alternative allele frequency (AF) in the population, whereas there is a gradually decreasing number of deletions at progressively higher AF. Over half of the insertions also occur at a low AF, but there is a sizeable number of insertions with very high AF or even becomes fixated (AF=1) in the population. As been reported previously^12^, these high AF insertions are likely to represent defects and/or rare alleles in the reference human genome. Based on the Hardy-Weinberg Equilibrium (HWE) test, we removed 2,868 (14%) SVs that are inconsistent with population genetics expectations. The removed SVs chiefly come from the unexpected AF peak at 0.5 (dashed lines in **Figure 5a**). 79% of these HWE-failed SVs are within TRs, which are likely to have higher mutation rates and be more variable in the population^33,34^. SVs that showed more genotyping errors in the discovery samples were more likely to fail the HWE test (**Table S3**). For example, while just 9% of the SVs with no genotyping errors failed our HWE test 40% of the SVs with two genotyping errors in our discovery samples failed our HWE test.

**Figure 5.**
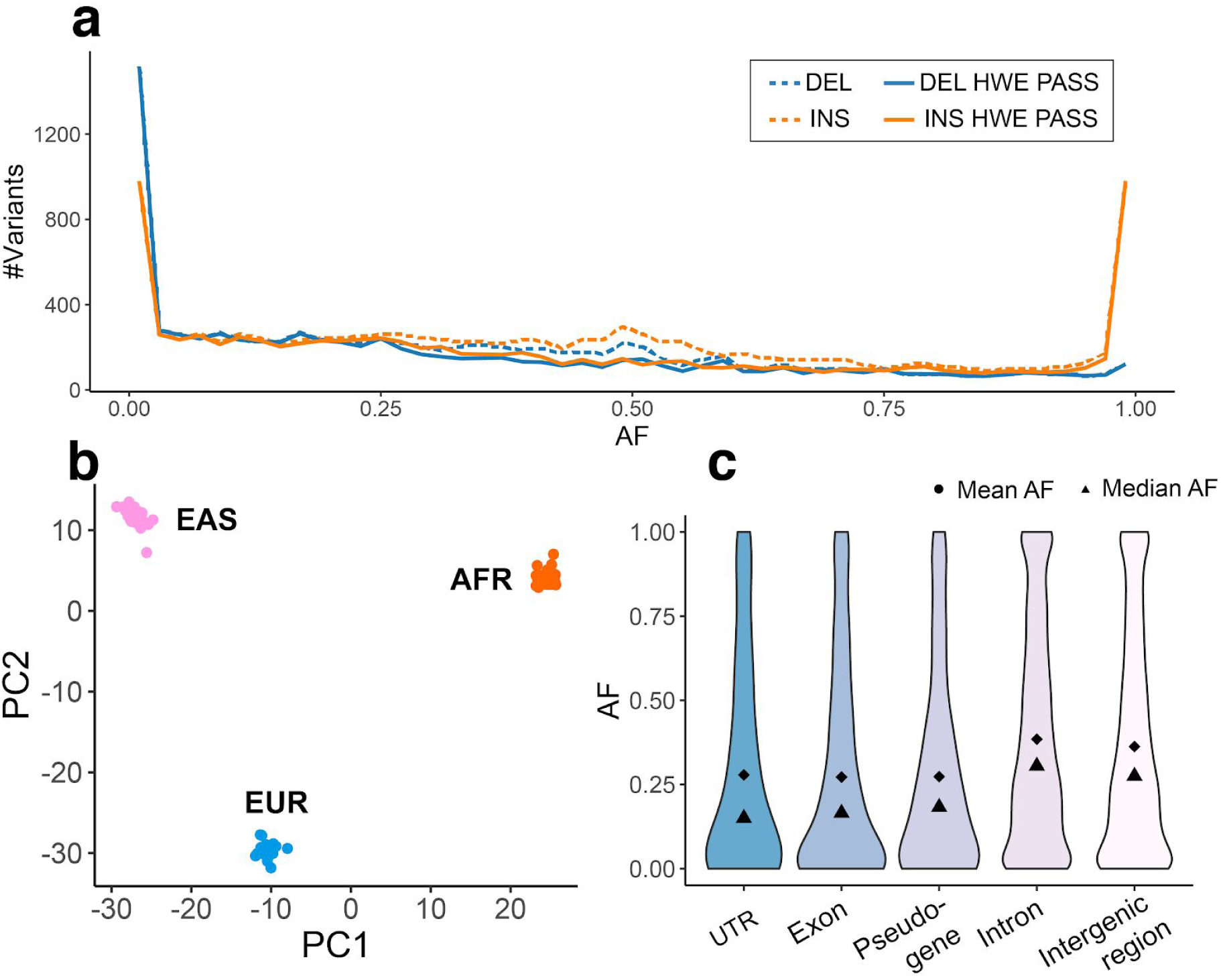
Population-scale genotyping and function annotation of LRGT SVs. (a) The AF distribution of LRGT SVs in the Polaris 100-individual population (b) PCA biplot of individuals in the population, based on genotypes of HWE-passing SVs. (c) The AF distribution of HWE-passing SVs in different functional elements. SV count: 191 in UTRs, 554 in exons, 420 in pseudogenes, 9,542 in introns and 6,603 in intergenic regions.

Because these samples are derived from different populations, our HWE test can be overly conservative, although only 962 (5%) of LRGT SVs have significantly different AFs between populations as measured by the test of their Fixation Index (F_st_)^35^. In the principal component analysis (PCA) of the HWE-passing SVs, the samples are clearly clustered by populations (**Figure 5b**). Interestingly, in PCA of the HWE-failed SVs, the samples cluster by population (**Figure S5**), too, indicating that some SVs could fail our HWE test because of population substructure rather than poor genotyping performance. Genotyping more samples in each of the three populations will allow better assessment of the genotyping accuracy without the confounding factor of subpopulations that could lead to erroneous HWE deviations.

The population AF can reveal information about the potential functional impact of SVs on the basis of signals of selective pressure. By checking the AFs for SVs in different genomic elements, we found that SVs within exons, pseudogenes and untranslated regions (UTRs) of coding sequences, in general, have lower AFs than those in intronic and intergenic regions. SVs in introns and intergenic regions have more uniform AF distributions compared to the more extreme AFs in functional elements (UTRs, exons) (**Figure 5c**). All these suggest a purifying selection against SVs with potentially functional consequences^25^. Common SVs are more depleted in functional regions than rare SVs, although we do see a few common SVs within exons of genes including *TP73* (AF=0.09, tumor suppressor gene), *FAM110D* (AF=0.60, functions to be clarified, possibly related with cell cycle) and *OVGP1* (AF=0.18, related to fertilization and early embryo development). As the three discovery samples are likely healthy individuals, and these SVs are found at a high frequency in the population, we expect them unlikely to have functional significance.

We also observed 17 exonic insertions fixated (AF=1) in the population (**Table S4**). Since these insertions are present and homozygous in all 100 genotyped individuals, the reference sequence reflects either rare deletion or errors in GRCh38^36^. Specifically, the 1,638 bp exonic insertion in *UBE2QL1* was also reported at high frequency in two previous studies^37,38^. Particularly, a recent study by TOPMed^38^ reported this insertion in all 53,581 sequenced individuals from mixed ancestries. Applying Paragraph to population-scale data will give us a better understanding of common, population-specific, and rare variations and aid in efforts to build a better reference genome.

## Discussion

Here we introduce Paragraph, an accurate graph-based SV genotyper for short-read sequencing data. Using SVs discovered from high-quality long read sequencing data of three individuals, we demonstrate that Paragraph achieves substantially higher recall (0.84 for deletions and 0.88 for insertions) compared to three commonly used genotyping methods (highest recall at 0.76 for deletions across the genome) and three commonly used *de novo* SV callers (highest recall of 0.64 for deletions). Of particular note, Paragraph and Manta were the only two methods that worked for both deletions and insertions and based on our test data Paragraph achieved substantially higher recall for insertions compared to Manta (0.88 vs 0.35).

As highlighted above, a particular strength of Paragraph is the ability to genotype both deletions and insertions genome-wide, including those within complicated regions. While we expect that there are as many insertions as there are deletions in the human population, the majority of the commonly used methods either do not work for insertions or perform poorly with the inserted sequence. In particular, because insertions are poorly called by *de novo* variant callers from short reads. Currently the most effective method to identify insertions is through discovery with long reads. Once a reference database of insertions is constructed, they can then be genotyped with high accuracy in the population using Paragraph. We expect this will be especially helpful to genotype clinically relevant variants as well as to assess variants of unknown significance (VUS) by accurately calculating AFs in healthy and diseased individuals.

Existing population reference databases for SVs may include many variants that are incorrectly represented. Since errors in the breakpoints may be a limitation for population-scaled SV genotyping, we have quantified the genotyping performance of Paragraph and its correlation with breakpoint accuracy (**Figure 3**). Our analysis shows that Paragraph can generally tolerate breakpoint deviation of up to 10 base pairs in most genomic contexts, although the performance suffers as the breakpoints deviate by more bases. Undoubtedly, recent advances in long read accuracy will lead to more accurate SV reference databases and thus better performance for Paragraph as a population genotyper.

Paragraph works by aligning and genotyping reads on a local sequence graph constructed for each targeted SV. This approach is different from other proposed and most existing graph methods that create a single whole-genome graph and align all reads to this large graph^18,39^. A whole-genome graph may be able to rescue reads from novel insertions that are misaligned to other parts of the genome in the original linear reference, however, the computational cost of building such a graph and performing alignment against this graph is very high. Adding variants to a whole-genome graph is also a very involved process that typically requires all reads to be realigned. Conversely, the local graph approach applied in Paragraph is not computationally intensive and can easily be adapted into existing secondary analysis pipelines. The local graph approach utilized by Paragraph also scales well to population-level studies where large sets of variants identified from different resources can be genotyped rapidly (e.g. 1,000 SVs can be genotyped in one sample in 15 minutes with a single thread) and accurately in many samples.

In this study, we demonstrated that Paragraph can accurately genotype single SVs that are not confounded by the presence of nearby SVs (**Table 1, Table S2**). Though, of the SVs identified in these three samples, almost half (48%) occurred in the presence of one or more different SVs. The current version of Paragraph only genotypes one SV per locus though we are actively working on the algorithm to consider and test the ability to annotate overlapping SVs and genotype them simultaneously. In addition, it will be equally important to create a more complete catalog of SVs in these highly variable loci so that the entire complexity can be encoded into the graph.

The primary use case for Paragraph will be to allow investigators to genotype previously identified variants with high accuracy. This could be applied to genotype known, medically relevant SVs in precision medicine initiatives or to genotype SVs from a reference catalog for more accurate assessment in a population or association study. Importantly, the catalog of both medically important SVs and population-discovered SVs will continue to evolve over time and Paragraph will allow scientists to genotype these newly-identified variants in historical sequence data. Certainly, the variant calls for both small (single sample) and large (population-level) sequencing studies can continue to improve as our knowledge of population-wide variation becomes more comprehensive and accurate.

## Conclusions

Paragraph is an accurate SV genotyper for short-read sequencing data that scales to hundreds or thousands of samples. Paragraph implements a unified genotyper that works for both insertions and deletions, independent of the method by which the SVs were discovered. Thus, Paragraph is a powerful tool for studying the SV landscape in populations, human or otherwise, in addition to analyzing SVs for clinical genomic sequencing applications.

## Methods

### Graph construction

In a sequence graph, each node represents a sequence that is at least one nucleotide long and directed edges define how the node sequences can be connected together to form complete haplotypes. Labels on edges are used to identify individual alleles or haplotypes through the graph. Each path represents an allele, either the reference allele, or one of the alternative alleles. Paragraph currently supports three types of SV graphs: deletion, insertion, and blockwise sequence swaps. Since we are only interested in read support around SV breakpoints, any node corresponding to a very long nucleotide sequence (typically longer than two times the average read length) is replaced with two shorter nodes with sequences around the breakpoints.

### Graph alignment

Paragraph extracts reads, as well as their mates (for paired-end reads), from the flanking region of each targeted SV in a Binary Alignment Map (BAM) or CRAM file. The default target region is one read length upstream of the variant starting position to one read length downstream of the variant ending position, although this can be adjusted at runtime. The extracted reads are realigned to the pre-constructed sequence graph using a graph-aware version of a Farrar’s Striped Smith-Waterman alignment algorithm implemented in GSSW library^40^ v0.1.4. In the current implementation, read pair information is not used in alignment or genotyping. The algorithm extends the recurrence relation and the corresponding dynamic programming score matrices across junctions in the graph. For each node, edge, and graph path, alignment statistics such as mismatch rates and graph alignment scores are generated.

Only uniquely mapped reads, meaning reads aligned to only one graph location with the best alignment score, are used to genotype breakpoints. Reads used in genotyping must also contain at least one kmer that is unique in the graph. Paragraph considers a read as supporting a node if its alignment overlaps the node with a minimum number of bases (by default 10% of the read length or the length of the node, whichever is smaller). Similarly, for a read to support an edge between a pair of nodes means its alignment path contains the edge and supports both nodes under the above criteria.

### Breakpoint genotyping

A breakpoint occurs in the sequence graph when a node has more than one connected edges. Considering a breakpoint with a set of reads with a total read count *R* and two connecting edges representing haplotype *h*_1_ and *h*_2_. We define the read count of haplotype *h*_1_ as *R*_*h*1_ and haplotype *h*_2_ as *R*_*h*2_. The remaining reads in *R* that are mapped to neither haplotype are denoted as *R*_≠*h*1,*h*2_.

The likelihood of observing the given set of reads with the underlying breakpoint genotype *G*_*h*1/*h*2_ can be represented as:

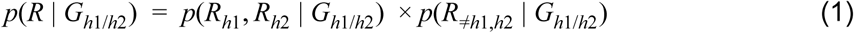

We assume that the count of the reads for a breakpoint on the sequence graph follows a Poisson-distribution with parameter λ. With an average read length *l*, an average sequencing depth *d*, and the minimal overlap of *m* bases (default: 10% of the read length *l*) for the criteria of a read supporting a node, the Poisson parameter can be estimated as

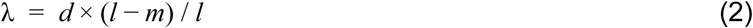

When assuming the haplotype fractions (expected fraction of reads for each haplotype when the underlying genotype is heterozygous) of *h*_1_ and *h*_2_ are μ_*h*1_ and μ_*h*2_, the likelihood under a certain genotype, *p*(*R*_*h*1_, *R*_*h*2_ | *G*_*h*1/*h*2_), or the first term in equation (1), can be estimated from the density function *dpois*() of the underlying Poisson distribution:

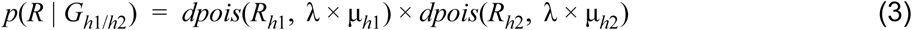

If *h*_1_ and *h*_2_ are the same haplotypes, the likelihood calculation is simplified as:

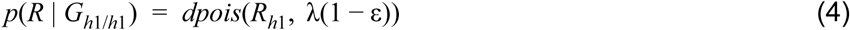

where ε is the error rate of observing reads supporting neither *h*_1_ nor *h*_2_ given the underlying genotype *G*_*h*1/*h*2_. Similarly, the error likelihood, *p*(*R*_≠*h*1,*h*2_ | *G*_*h*1/*h*2_), or the second term in equation (1), can be calculated as

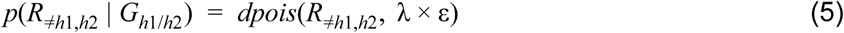

Finally, the likelihood of observing genotype *G*_*h*1/*h*2_ under the observed reads *R* can be estimated under a Bayesian framework

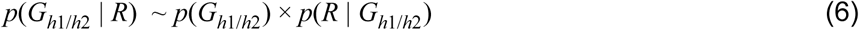

The prior *P* (*G*_*h*1/*h*2_) can be pre-defined or calculated using a helper script in Paragraph repository that uses the Expectation-Maximization algorithm to estimate genotype-likelihood based allele frequencies under Hardy-Weinberg Equilibrium across a population^41^.

### SV genotyping

1. We perform a series of tests for the confidence of breakpoint genotypes. For a breakpoint to be labeled as “passing”, it must meet all of the following criteria:
2. It has more than one read aligned, regardless of which allele the reads were aligned to
3. The breakpoint depth is not significantly high or low compared to the genomic average (p-value is at least 0.01 on a two-sided Z-test)
4. The Phred-scaled score of its genotyping quality (derived from genotype likelihoods) is at least 10.
5. Based on the reads aligned to the breakpoint, regardless of alleles, the Phred-scaled p-value from FisherStrand^42^ test is at least 30.

If a breakpoint fails one or more of the above tests, it will be labeled as a “failing” breakpoint. Based on the test results of the two breakpoints, we then derive the SV genotype using the following decision tree:

1. If two breakpoints are passing:
  a. If they have the same genotype, use this genotype as the SV genotype
  b. If they have different genotypes, pool reads from these two breakpoints and perform the steps in **Breakpoint genotyping** section again using the pooled reads. Use the genotype calculated from the pooled reads as the SV genotype.
2. If one breakpoint is passing and the other one is failing:
  a. use the genotype from the passing breakpoint as the SV genotype.
3. If two breakpoints are failing:
  a. If the two breakpoints have the same genotype, use this genotype as the SV genotype
  b. If two breakpoints have different genotypes, follow the steps in 1b.

Note that for 1b and 2b, as we pool reads from two breakpoints together, the depth parameter *d* in equation (2) needs to be doubled, and reads that span two breakpoints will be counted twice. We also set a filter label for the SV after this decision tree, and this filter will be labeled as passing only when the SV is genotyped through decision tree 1a. SVs that fail the passing criteria 1 and 2 for any one of its breakpoints were considered as reference genotypes in the evaluation of Paragraph in the main text.

### Sequence data

The CCS data for NA12878 (HG001), NA24385 (HG002) and NA24631 (HG005) are available at the GiaB FTP (ftp://ftp.ncbi.nlm.nih.gov/giab/ftp/data/). These samples were sequenced to an approximate 30x depth with an average read length of 11 kb on the PacBio Sequel system. We re-aligned reads to the most recent human genome assembly, GRCh38, using pbmm2 v1.0.0 (https://github.com/PacificBiosciences/pbmm2). Pacbio CLR data of NA24385^11^ were sequenced to 50x coverage on a PacBio RS II platform, and reads were aligned to GRCh38 using NGMLR^10^ v0.2.7.

To test the performance of the methods on short-read data, we utilized three matching samples that were sequenced using TruSeq PCR-free protocol on Illumina platforms with 150 bp paired-end reads: 35x (NA24385) on HiSeq X, 64x (NA12878) and 48x (NA24631) on NovaSeq 6000. Reads were mapped to GRCh38 using the Issac aligner^43^. To estimate the recall of Paragraph in samples of lower depth, we downsampled the 35x NA24385 data to different depths using samtools^44^. To estimate the recall of Paragraph in 100bp and 75bp reads, we trimmed the 150bp reads from their 3’ end in the downsampled NA24385 data.

### Long read ground truth and performance evaluation

SVs were called from the CCS long read data of the three samples using PBSV v2.0.2 (https://github.com/PacificBiosciences/pbsv). When merging SVs across samples, we define deletions as “different” if their deleted sequences have less than 80% reciprocal overlap; we define insertions as “different” if their breakpoints are more than 150 bp apart, or their insertion sequences have less than 80% of matching bases when aligning against each other using Smith-Waterman algorithm. After merging, we obtained 41,186 unique SVs. From these unique SVs, we excluded 1,944 from chromosome X or Y, 53 SVs that had a failed genotype in one or more samples and 480 SVs where a nearby duplication was reported in at least one sample. In the remaining 38,709 unique SVs, 20,108 have no nearby SVs within 150bp upstream and downstream and these SVs were used as LRGT to test the performance of Paragraph and other methods.

For each method, we define a variant as a true positive (TP) if the LRGT data also has a call in the same sample and a false positive (FP) if the LRGT did not call a variant in that sample. For each genotyper, we estimate its recall as the count of its TPs divided by the count of alternative genotypes in LRGT. We calculate the precision of each method as its TPs divided by its TPs plus FPs. Variants identified by the *de novo* methods (Manta, Lumpy, and Delly) may not have the same reference coordinates or insertion sequences as the SVs in LRGT. To account for this we matched variants from *de novo* callers and SVs in LRGT using Illumina’s large-variant benchmarking tool, Wittyer (v0.3.1). Wittyer matches variants using centered-reciprocal overlap criteria, similar to Truvari (https://github.com/spiralgenetics/truvari) but has better support for different variant types and allow stratification for variant sizes. We set parameters in Wittyter as “--em simpleCounting --bpd 500 --pd 0.2”, which means for two matching variants, their breakpoint needs to be no more than 500 bp apart from each other and if they are deletions, their deleted sequences must have no less than 80% reciprocal overlap.

### Estimation of breakpoint deviation

From CLR NA24385, SVs were called using the long read SV caller, Sniffles^10^ with parameters “--report-seq -n -1” to report all supporting read names and insertion sequences. Additional default parameters require 10 or more supporting reads to report a call, and require variants to be at least 50 bp in length. Insertion calls were refined using the insertion refinement module of CrossStitch (https://github.com/schatzlab/crossstitch), which uses FalconSense, an open-source method originally developed for the Falcon assembler^45^ and is also used as the consensus module for Canu^46^.

We used a customized script to match calls between the CLR and LRGT SVs of NA24385. A deletion from the CLR data is considered to match a deletion in LRGT if their breakpoints are no more than 500 bp apart and their reciprocal overlap length is no less than 60% of their union length. An insertion from the CLR data is considered to match an insertion in LRGT if their breakpoints are no more than 500 bp apart. Base pair deviations between insertion sequences were calculated from the pairwise alignment method implemented the python module biopython^47^.

### Population genotyping and annotation

The 100 unrelated individuals from the Polaris sequencing resource (https://github.com/Illumina/Polaris) were sequenced using TruSeq PCR-free protocol on Illumina HiSeq X platforms with 150 bp paired-end reads. Each sample was sequenced at an approximate 30-fold coverage. We genotyped the LRGT SVs in each individual using Paragraph with default parameters.

For each SV, we used Fisher’s exact test to calculate its Hardy-Weinberg p-values^48^. SVs with p-value less than 0.0001 were considered as HWE-failed. We used dosage of HWE-passing SVs to run PCA, which means 0 for homozygous reference genotypes and missing genotypes, 1 for heterozygotes and 2 for homozygous alternative genotypes.

We used the annotation tracks from the UCSC Genome Browser to annotate SVs in LRGT. We define an SV as “within TR” if its reference sequence is completely within one or more TRF tracks. We categorized an SV as functional if it overlaps with one or more functional tracks. We used the ENCODE Exon and PseudoGene SupportV28 track for exons, IntronEst for introns and ENCFF824ZKD for UTRs. SVs that overlap with any functional tracks SVs that do not overlap with any of these tracks were annotated as intergenic.

## Acknowledgments

We thank Mitch Bekritsky and Camilla Colombo for their support in the sequence data, and Chi Kent Ho and Yinan Wan for their help in evaluating the genotyping performance of different methods.

## Availability of data and materials

Paragraph software is publicly available on https://github.com/Illumina/paragraph. All analysis in the manuscript was performed under version 2.3 deposited at DOI/10.5281/zenodo.3440238. The three Illumina sequenced samples will be available soon. SV calls from the three long read sequenced samples, and individual genotypes in Polaris cohort of these SVs, are available in Paragraph GitHub repository.

## Authors’ contributions

PK, SC, ED, RP, and FS developed the algorithms and wrote the source code. MAE, SC, FJS, and MCS designed the manuscript framework. SC performed the experiments, prepared the figures and wrote the manuscript. FJS, RMS, and MK prepared the test set on long read data. MAE, DRB, FJS, MCS, RMS, PK, and ED edited the manuscript. All authors read and approved the manuscript.

## Ethics approval and consent to participate

Not applicable.

## List of abbreviations

SV: structural variation
bp: base pair
TR: tandem repeat

