## Supplemental Tables and Figures for "Paragraph: A graph-based structural variant genotyper for short-read sequence data"

### Supplementary Figures and Tables

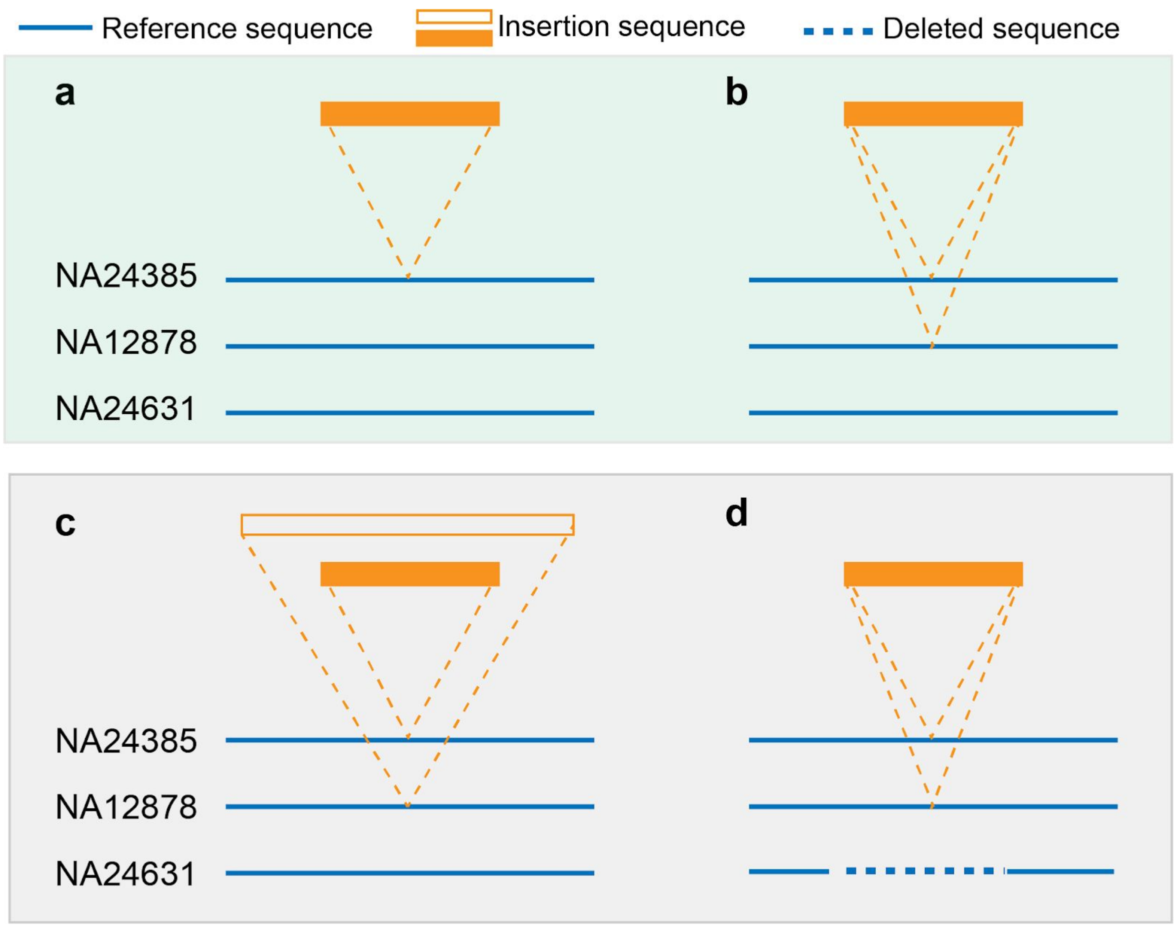


**Figure S1. Examples of singleton SVs and clustered SVs.** Variants that were called in only one sample, or called from more than one sample but have the same description, were labeled as singleton SVs and were included in our LRGT. Examples as (a) and (b) in the green box. Variants that were called in more than one sample with different descriptions were labeled as clustered SVs and were excluded from LRGT. Examples as (c) and (d) in the grey box.

(a) An insertion which is called in only one sample
(b) An insertion which is called in two samples and has the same insertion sequence
(c) Two of the three samples have an insertion called at a similar position but the insertion sequences are different between two samples
(d) Two of the three samples have the same insertion called but the third sample has a deletion that overlaps with this insertion

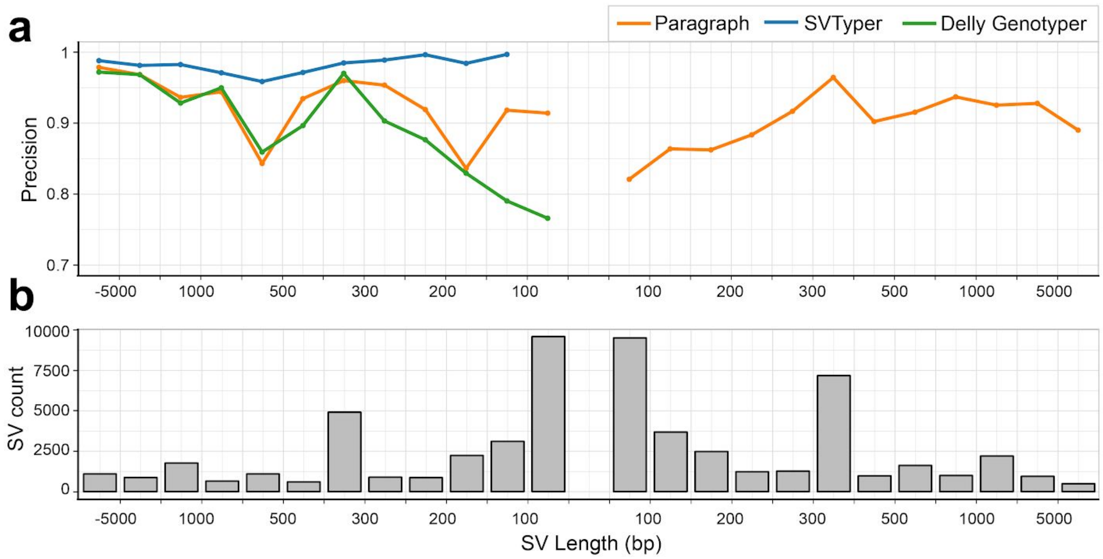


**Figure S2. Estimated precision of different genotypers, partitioned by SV length**. Precision was estimated on short-read sequencing data of the three samples using LRGT as the truth set. Colored lines in (a) show precision of different genotypers; Solid grey bars in (b) represent the count of SVs tested in each size range.

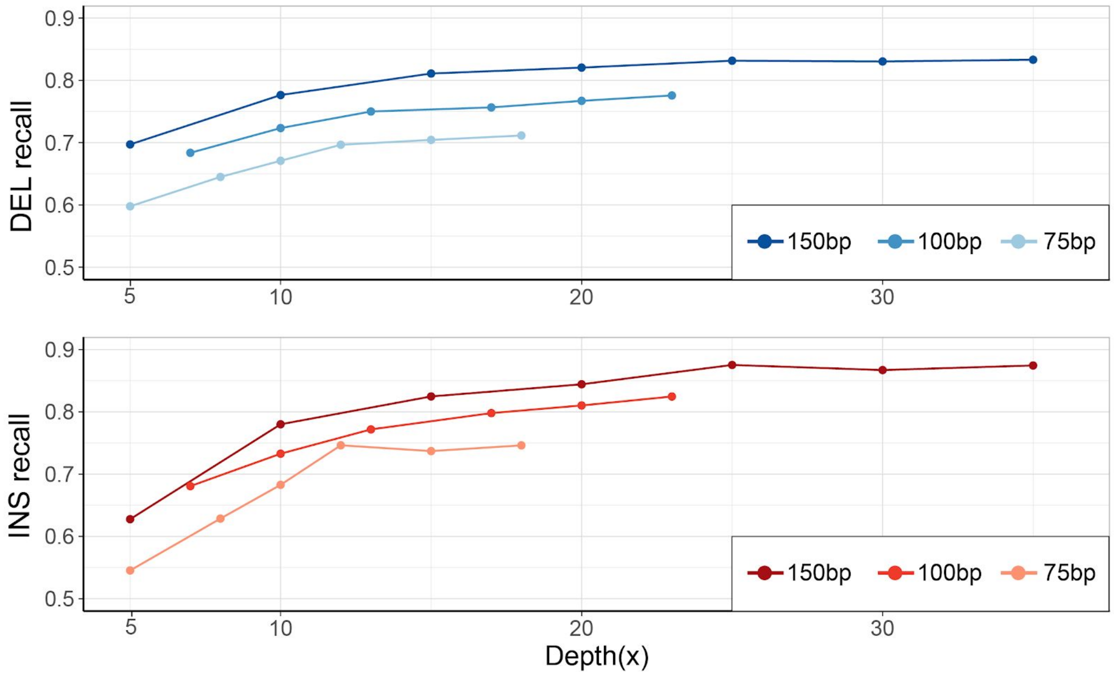


**Figure S3. Recall under different read lengths and depths.** Paragraph recall was measured on short read sequencing data of NA24385 against LRGT. Test data under different lengths and sequencing depths was generated by downsampling and trimming on the 150bp paired-end reads of NA24385 short read data.

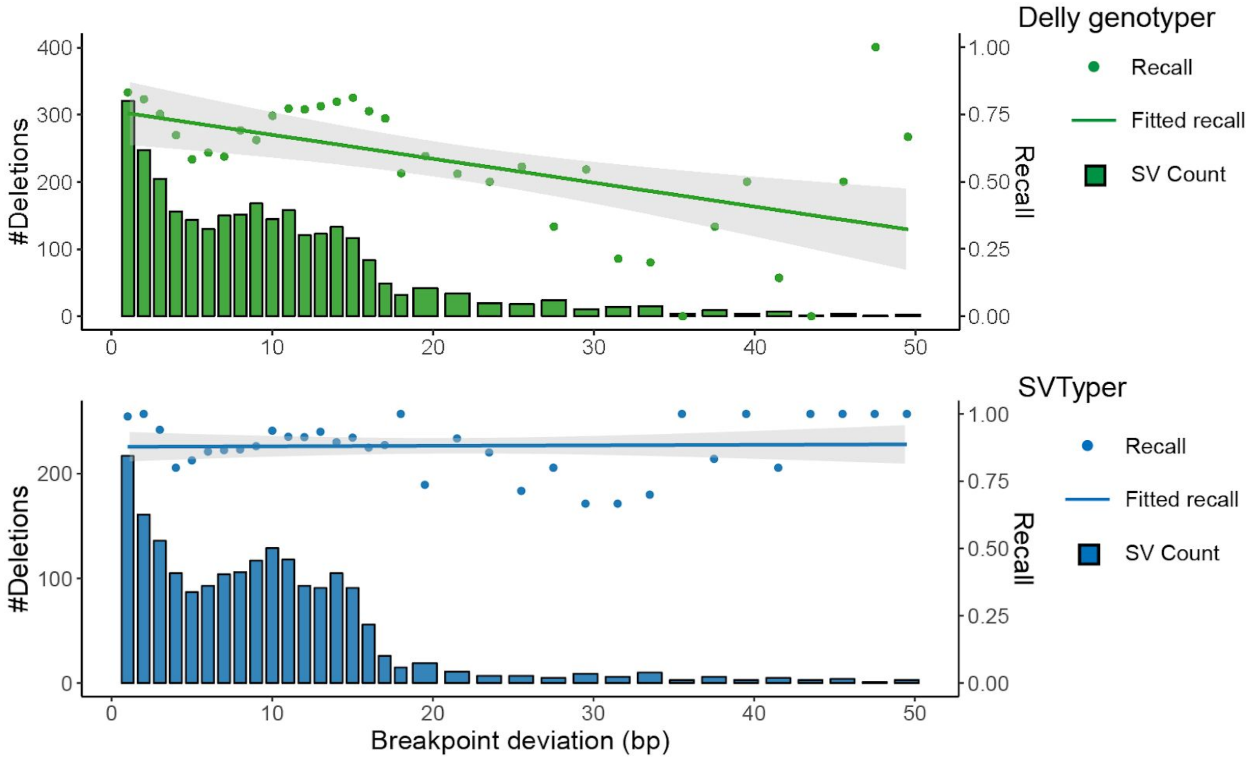


**Figure S4. The impact of recall when tested SVs include errors in their breakpoints.** Follow the same workflow as in **Figure 3**, Delly Genotyper and SVTyper recall were estimated using CLR calls as genotyping input and their TPs in LRGT as the ground truth, respectively.

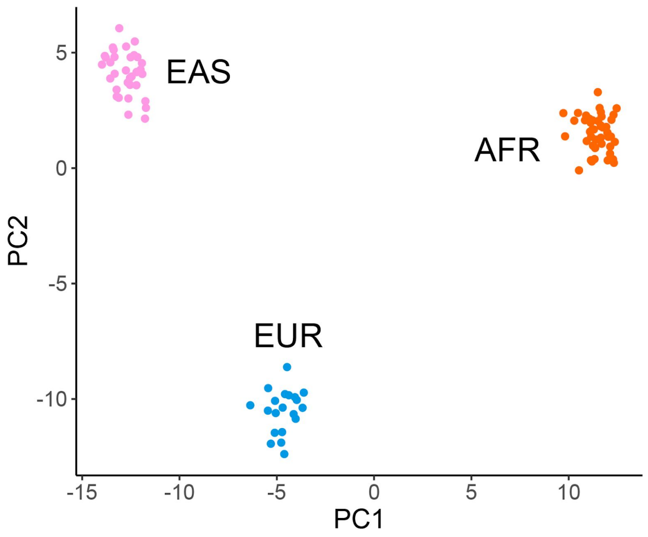

**Figure S5. PCA biplot of the Polaris population using only HWE-failed SVs.** Genotypes of HWE-failed SVs were used as the input for PCA.

| | Deletion | | | Insertion |
|---|---|---|---|---|
| **Genotyper** | Paragraph | Delly Genotyper | SVTyper (100+bp) | Paragraph |
| **Concordance** | 0.81 | 0.73 | 0.73 | 0.72 |

**Table S1. Estimated genotype concordance of genotypers.** The concordance was estimated on short-read data of the three samples, with LRGT genotypes as the ground truth. A genotype is considered as concordant only if the genotype from short-read genotyper output is exactly the same as in LRGT.

| | Deletion | | | | | | Insertion | |
|---|---|---|---|---|---|---|---|---|
| **Type** | **Paragraph** | **Delly Genotyper** | **SVTyper (100+ bp)** | **Manta** | **Delly** | **Lumpy (100+ bp)** | **Paragraph** | **Manta** |
| #Tested TPs | 10,873 | 10,873 | 6,900 | 10,919 | 10,919 | 6,900 | 15,996 | 15,996 |
| Recall | 0.71 | 0.65 | 0.30 | 0.19 | 0.20 | 0.15 | 0.79 | 0.13 |
| #Tested FPs | 12,272 | 12,272 | 7,602 | - | - | - | 16,662 | - |
| Precision | 0.58 | 0.52 | 0.69 | - | - | - | 0.54 | - |
| F-score | 0.64 | 0.58 | 0.42 | - | - | - | 0.64 | - |

**Table S2. Performance of different methods in clustered SVs.** Genotyping/calling was evaluated on short read data of the same three discovery samples as described in Table 1.

| | HWE status | | |
|---|---|---|---|
| **#SVs with genotyping errors** | PASS | FAIL | **%HWE-failed SVs** |
| 0 | 13,283 | 1,335 | 9% |

| | | | |
|---|---|---|---|
| 1 | 2,487 | 693 | 22% |
| 2 | 1,127 | 766 | 40% |
| 3 | 378 | 39 | 9% |

**Table S3. SVs with genotyping errors in the three discovery samples.** For each LRGT SV in each sample, if the presence/absence of this SV is different between the Paragraph genotype and the LRGT genotype, it is counted as one genotyping error. As there are three samples in LRGT, the maximum genotyping error for an SV is three. An SV fails HWE if its HWE p-value in the Polaris population is smaller than 0.0001.

| Chromosome | Start | LRGT ID | RefSeq gene |
|---|---|---|---|
| chr1 | 41382200 | HG002_pbsv.INS.1008 | *FOX06* |
| chr1 | 247937455 | HG002_pbsv.INS.4079 | *OR2L13* |
| chr11 | 47557553 | HG002_pbsv.INS.33173 | *RN7SL652P* |
| chr12 | 123389131 | HG002_pbsv.INS.36631 | *KMT5A* |
| chr12 | 127634170 | HG002_pbsv.INS.36777 | *LINC02411* |
| chr13 | 36794664 | HG002_pbsv.INS.37593 | *ARL2BPP3* |
| chr15 | 90871538 | HG002_pbsv.INS.41892 | *FURIN* |
| chr17 | 82443764 | NA12878_pbsv.INS.44159 | *CYBC1* |
| chr18 | 58863528 | HG002_pbsv.INS.46767 | *ZNF532* |
| chr19 | 14090084 | HG002_pbsv.INS.47894 | *SAMD1* |
| chr2 | 240527763 | HG002_pbsv.INS.7994 | *ANKMY1* |
| chr20 | 35226721 | HG002_pbsv.INS.49926 | *EDEM2* |
| chr22 | 20268314 | HG002_pbsv.INS.52069 | *RTN4R* |
| chr5 | 6448622 | HG002_pbsv.INS.15266 | *UBE2QL1* |
| chr6 | 168029510 | HG002_pbsv.INS.20934 | *KIF25* |

| | | | |
|---|---|---|---|
| chr7 | 73578776 | HG002_pbsv.INS.22659 | *TBL2* |
| chr8 | 124309834 | NA12878_pbsv.INS.25530 | *TMEM65* |

**Table S4. List of fixated exonic SVs.** SVs in the table were genotyped as homozygous alternative in all 100 individuals in the Polaris population, and occur in the exons of affected genes.
